# Evaluating the Effectiveness of Data Reduction Techniques in QTL Mapping

**DOI:** 10.1101/2025.08.29.673132

**Authors:** Caroline Keller, Celine Caseys, Daniel J. Kliebenstein

## Abstract

Data reduction methods are frequently employed in large genomics and phenomics studies to extract core patterns, reduce dimensionality, and alleviate multiple testing effects. Principal component analysis (PCA), in particular, identifies the components that capture the most variance within omics datasets. While data reduction can simplify complex datasets, it remains unclear how the use of PCA impacts downstream analyses such as quantitative trait loci (QTL) or genome-wide association (GWA) approaches and their biological interpretation. In QTL studies, an alternative to data reduction is the use of post-hoc data summarization approaches, such as hotspot analysis, which involves mapping individual traits and consolidating results based on shared genomic locations. To evaluate how different analytical approaches may alter the biological insights derived from multi-dimensional QTL datasets, we compared individual trait hotspots with PCA-based QTL mapping using transcriptomic and metabolomic data from a structured recombinant inbred line population. Interestingly, these two approaches identified different genomic regions and genetic architectures. These findings suggest that mapping PCA-reduced data does not merely streamline analyses but may generate a fundamentally different view of the underlying genetic architecture compared to individual trait mapping and hotspot analysis. Thus, the use of PCA and other data reduction techniques prior to QTL or GWAS mapping should be carefully considered to ensure alignment with the specific biological question being addressed.

## Introduction

Modern omics technologies such as transcriptomics, metabolomics, and phenomics generate massive datasets that are challenging to wrangle and whose size complicates interpretation. When faced with these massive datasets, data reduction approaches are commonly performed to simplify computational intricacy and resources. An additional benefit of data reduction is the enhanced statistical power due to a lower number of statistical tests and a lower false discovery rate when the data is consolidated (Jollife & Cadima, 2016). This is often associated with the assumption that data reduction helps clarify data interpretation and visualization by simplifying the data to its key messages. However, it is not always clear whether data decomposition aids in analysis, retains most of the biological information, and is empirically comparable to the original dataset.

One of the most common data decomposition approaches is principal component analysis (PCA), which reduces the dimensionality of multivariate datasets into principal component vectors. Depending on the specific approach, PCA computes the maximal correlation or covariance values across all the data points by estimating linear combinations of variables and extracting the resulting vector. The process is then repeated with the residuals to identify additional PCs associated with decreasing proportions of variance (Camargo, 2022). This approach is useful in natural variation or quantitative genetics omics studies, as the original factors/traits contributing to the PCs with the highest variance can be identified, and each vector can have a discrete mathematical value to use as a composite trait for genetic mapping (Vieira, 2012). However, the use of the correlational or covariance matrix for PCA will alter the underlying question by focusing either on the vectors that influence the largest percentage of variation (correlation) or the largest magnitude of variation (covariance). This suggests that these approaches might identify different genetic loci.

The alternative to data decomposition approaches is post-hoc summary methods wherein all the individual traits (transcripts, metabolites, or phenotypes) are mapped one-by-one (Breitling et al., 2008). The position of each QTL/trait combination is then used to find the genomic locations affecting the greatest number of individual traits. In this method, the loci of interest are defined using a permutation approach that determines how many traits are expected to randomly associate with a specific genomic region. Hotspot analysis can identify functional loci that offer insights into altered molecular pathways, enabling the formulation of hypotheses and the discovery of their causal basis (Francisco, Kliebenstein, Rodríguez, & Soengas, 1871; Li et al., 2016; Rowe, Hansen, Halkier, & Kliebenstein, 2008). However, this methodology, focusing on the loci with the greatest number of associated traits, can create problems. For example, permutation thresholding may fail to detect QTLs that influence specific pathways, as their limited and focused effects may not involve enough individual traits to surpass the hotspot significance threshold (Kliebenstein et al., 2006).

To evaluate how different normalization methods impact the interpretation of omics QTL datasets, we compared three methods using two different strategies: use of absolute transcript/metabolite values or log2 normalized values. The first method was to conduct QTL mapping on all individual traits and summarize the results using the standard hotspot approach, which counts the number of traits (transcripts or metabolites) influenced by genetic variation at each locus. The second method applied PCA to the correlation matrix to generate components that represent the percentage of variation within each individual trait. The third method applied PCA to the covariance matrix, generating components that explain the most variable individual traits. The inclusion of these three normalization approaches allows for an assessment of how sensitive each method is to encompass typical normalization methods used for omics datasets.

To compare the performance of PCA and post-hoc summary approaches, we performed QTL mapping on metabolomics and transcriptomics datasets from a bi-parental population of *Arabidopsis thaliana*. The recombinant inbred lines (RILs) of Bay-0 x Sha were originally developed and genotyped by Loudet et al. (2002) (http://www.inra.fr/qtlat; Loudet et al., 2002; West et al., 2007; Rowe et al., 2008). This RIL population is beneficial for this comparison because it has extensive recombination, and all individuals are homozygous with minimal population structure. This allows replicated omics experiments in which the environment is controlled, and technical factors are randomized as much as possible. These experimental conditions should maximize the PCA’s ability to overlook environmental components and attempt to capture the genetic information.

In this study, we evaluated how QTL mapping based on principal components (PCs) of metabolomic and transcriptomic datasets compares to mapping individual traits and post-hoc hotspot analyses. The genomic position and loadings of the PC-associated QTLs was compared to post-hoc hotspot summaries consolidated from the QTLs of individual transcripts and metabolites. We extended the analysis to map QTLs up to PC 211 to determine the point at which no mappable information could be identified. While all approaches ultimately identified similar regions of the genome influencing overall phenotypic variation, the PC vector mapping and post-hoc hotspot summary approaches gave different impressions of the genetic architecture for both metabolomic and transcriptomic datasets. Further, this discrepancy was consistent regardless of data normalization or PCA strategies. Finally, QTLs were mappable well beyond the first 50 PCs, indicating that data reduction approaches in QTL or GWA mapping are not readily interchangeable with post-hoc summary approaches of individual trait analyses.

## Materials and Methods

### Metabolomic and Transcriptomic data for Bay-0 x Sha RILs

Publicly available metabolomic and transcriptomic datasets for 211 *A. thaliana* recombinant inbred lines (RILs) from the Bay-0 x Sha mapping population were used to compare the effectiveness of using PCA to summarize metabolite QTLs, capture metabolite variation, locate hotspots, and significant loci (Rowe, Hansen, Halkier, & Kliebenstein, 2008; Loudet, Chaillou, Camilleri, Bouchez, & Daniel-Vedele, 2002). The metabolomics dataset reports primary metabolites quantified by gas chromatography coupled with mass spectrometry (GC-MS) (Fiehn, Wohlgemuth, & Scholz, 2005; Loudet et al., 2002; Meyer et al., 2007; Nikiforova et al., 2005; Roessner et al., 2001). The transcriptomics dataset resulted from a microarray experiment with two conditions, either a control silwet L77 (SW) or 0.3mM salicylic acid (SA) (West et al., 2007; http://www.arabidopsis.org). Overall, data for a total of 371 metabolites and 22,810 transcripts were used for PCA and QTL mapping.

### QTL Mapping

The Bay-0 x Sha population was originally developed and genotyped by (Loudet et al., 2002) http://www.inra.fr/qtlat) and additional markers were added by transcriptome genotyping (West et al., 2006). The combined marker set was used to conduct QTL mapping using R/qtl2 (K. Broman et al., 2018; K. W. Broman et al., 2019) on the individual metabolites or transcripts, in addition to their respective derived PCs. QTL mapping of both original and reduced data allows for the direct comparison between the hotspot summary (post-QTL consolidation) and PC vector mapping (pre-QTL data reduction) approaches. For each dataset, files containing genotypes, phenotypes (metabolites, transcripts, or PCs), a genetic map (.gmap), and an instruction file (.yaml) were created. The original genetic map was utilized, and no new pseudo-markers were inserted (step=0) for the marker/pseudo-marker map (Rowe, Hansen, Halkier, & Kliebenstein, 2008). The genome probabilities were calculated with an error probability of 0.2%. The genome scan function was utilized with standard parameters, and the find peaks function was used with a threshold = 2, peakdrop = 1.5, and drop = 1.5. A sliding window approach (5cM windows with 2.5cM steps) was applied to the logarithm of the odds (LOD) peak output to summarize the number of metabolite or transcriptomic QTLs across all 5 chromosomes.

In addition to the LOD, the scan1coef function was used to calculate the effect size of each genetic marker and the total variance they captured across the PC traits. The proportion of variance for each PC vector was multiplied by the effect size of each PC vector at each marker to generate the weighted effect. The weighted effects were then summed across all vectors at each maker to generate the total variance attributable to this position and plotted by position on each chromosome.

### Permutation Analysis

To determine significance thresholds for the hotspot summaries, we applied a permutation test to assess the probability of random associations (Churchill & Doerge, 1994). Within each of 1000 random permutations of the genetic markers, the sliding window with the maximal number of metabolites or transcripts, random QTL associations were identified. At a 5% chance of random associations (alpha = 0.05), the threshold for a significant metabolite hotspot was 18, while the threshold for a significant transcript hotspot was 66. Hotspots hosting more QTLs than these values are reported.

### Statistical Analysis

All statistical analysis and visualization were performed using R Statistical Software (v4.2.2; R Core Team 2022).

To account for and reduce metabolite and transcript heteroscedasticity, which is commonly introduced due to uninduced biological variation (Sun & Xia, 2024), a Log2 transformation was performed on both datasets. Prior to the log2 transformation, the smallest non-zero value for each dataset was added to traits, technically removing zero values. Although the left-skew and heteroscedasticity were not completely resolved, the metabolite data were approaching a Gaussian distribution. After log 2 transformation, the transcript data followed a Gaussian distribution.

PCA was performed with the prcomp function in R on 211 metabolites (Rowe et al., 2008) and two transcriptomics datasets (West et al., 2007). The PCA was run using the covariance prcomp(data, center = TRUE, scale. = FALSE) and the correlation prcomp(data, center = TRUE, scale. = TRUE) matrix. All PCs for each dataset were used for QTL mapping. Furthermore, to align with commonly applied approaches for omics data, all analyses were run using absolute (raw data) and Log2 normalization methods. The combination of these parameters created six different approaches to perform data analysis for each dataset.

Approach 1: Absolute Metabolite, Transcriptomics SA or SW data → QTL mapping and analysis

Approach 2: Absolute Metabolite, Transcriptomics SA or SW data → PCA Correlation Matrix → QTL mapping and analysis

Approach 3: Absolute Metabolite, Transcriptomics SA or SW data → PCA Covariance Matrix → QTL mapping and analysis

Approach 4: Log2 normalized Metabolite, Transcriptomics SA or SW data → QTL mapping and analysis

Approach 5: Log2 normalized Metabolite, Transcriptomics SA or SW data → PCA Correlation Matrix → QTL mapping and analysis

Approach 6: Log2 normalized Metabolite, Transcriptomics SA or SW data → PCA Covariance Matrix → QTL mapping and analysis

## Results

### QTL Hotspots associated with Metabolites

To understand the effects of data reduction on QTL mapping, we first focused on a metabolomics dataset that contains 371 metabolites measured across 211 RIL progeny using a randomized block design. Initially, we regenerated QTL mapping for all the individual 371 metabolites using the absolute and log2 normalized mean values per metabolite (Rowe et al., 2008) (Figure 2). This identified 778 QTLs associated with 243 metabolites located in 11 major hotspots as previously reported (Rowe et al., 2008) (Figure 2). The largest QTL hotspot, Met.I.80 locus, is associated with 57 unique metabolites around position 80cM on chromosome I (Figure 2, Supplemental SLW output tables). To assess the effects of data transformation prior to PCA on QTL mapping, a Log2 transformation was applied to normalize the data and reduce skew before mapping QTLs (Figure 2). The log2 transformed metabolite QTL mapping identified a nearly identical pattern with a total of 687 QTLs. The hotspots are in the same locations as the QTL mapping of absolute values. The largest locus, Met.I.80, is associated with 57 metabolites in the original dataset, and 53 metabolites in the log2 transformed dataset, with 21 metabolites identified across methods. Thus, the log2 transformation shifted individual metabolite QTLs but identified a largely consistent pattern of hotspots (Figure 2). This analysis showed that R/qtl2 mapping found the same hotspots as previously found using composite interval mapping with QTL cartographer (Rowe et al., 2008) and that the normalization approach minimally influenced the post-hoc summary approach.

### PCA Consolidation of Large Metabolite Datasets

To test how data reduction affects the QTL mapping of metabolites, both correlation and covariance-based PCA were applied to the average metabolite accumulation across the Bay-0 X Sha RILs. For each approach, QTL mapping was run on the 211 principal components using R/qtl2. Furthermore, to assess if the log2 transformation modified the PCA analysis, QTL mapping was also performed on the PCA of log2 transformation data. The hypothesis was that QTL mapping of log2 and correlation matrix of transformed PCs might perform better, as the transformation of the raw data would reduce heteroscedasticity inherent in biological data. This produced four different PCA and normalization data sets that assessed the problem with multiple approaches (Figure 1). Mapping the 211 PCs for the absolute metabolite values revealed 161 QTLs for covariance and 174 QTLs for the correlation datasets. Mapping the log2 normalized metabolite values found 132 QTLs for covariance and 146 QTLs for the correlation datasets (Supplemental SLW output tables). The four different PCA datasets identified different QTLs that were largely evenly distributed across the chromosomes. Interestingly, the PC QTL mapping approach did not track with the post-hoc hotspot summaries, and the first PCs failed to capture the largest hotspots (Figure 3A-3D).

**Figure 1:**
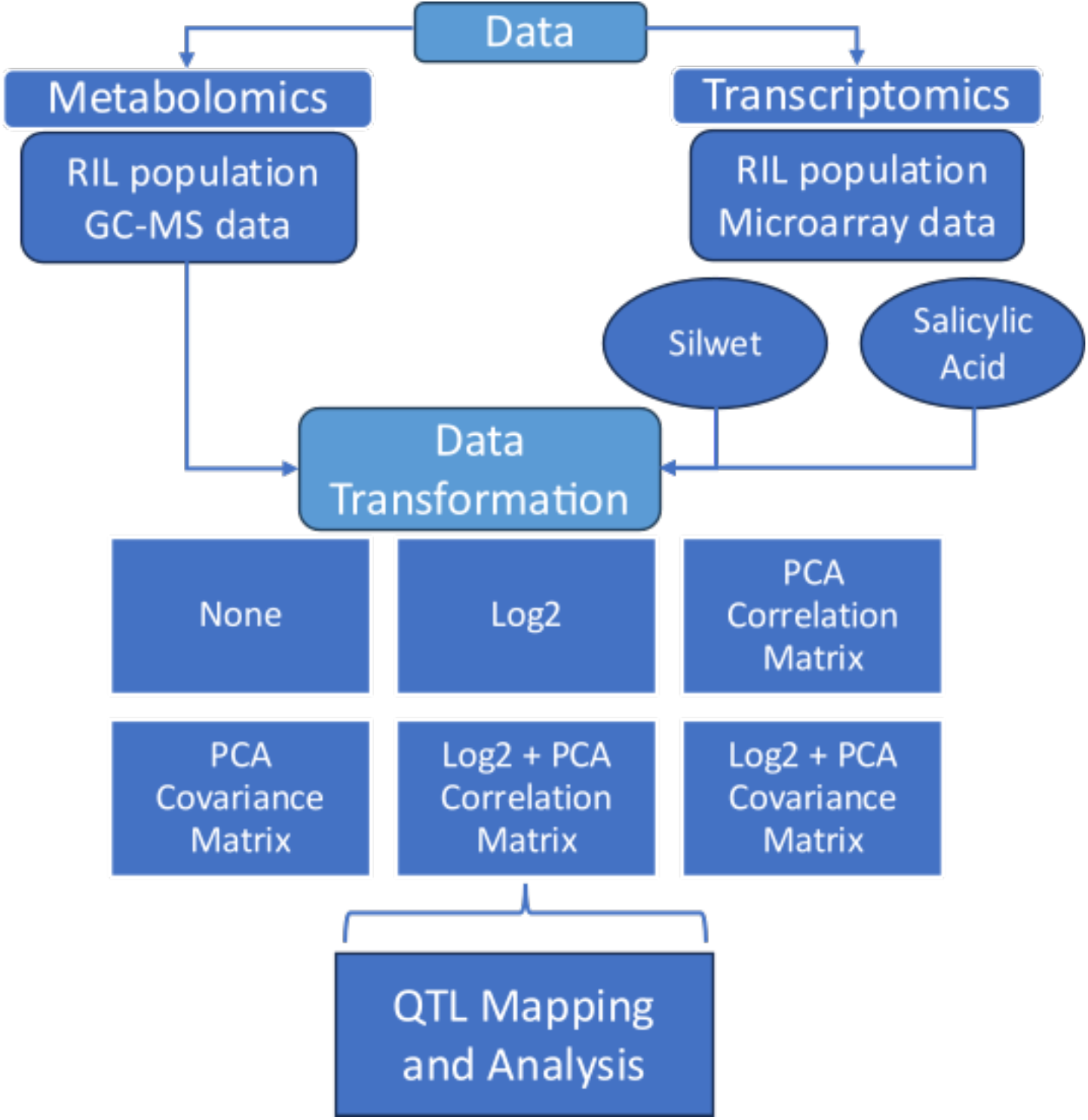
Protocol and data analysis flowchart. Metabolomics and Transcriptomics datasets from an *A. thaliana* RIL population were analyzed with R/qtl2. Each dataset was given no transformation, Log2 transformation, PCA with the correlation or covariance matrix, and a combination of the Log2 and PCA transformation. The six data transformation datasets were subsequently analyzed by QTL mapping.

One hypothesis was that the first PCs that capture most of the variance would identify the largest hotspots found in the individual trait analysis. This hypothesis was rejected as the largest post-hoc summary hotspots did not colocalize with the first PCs. The largest hotspots were typically found within the top 50 PCs and often found by multiple independent PCs (Figure 6 and Supplemental Figure 1 and 2). Additionally, we identified QTLs for the top 50 PCs, with additional QTLs being identified well beyond the top 100 PCs (Figure 6A-6D). Individual PCs identified multiple QTLs, and some genetic positions identified QTLs for multiple PCs. This was equally true for all four PCA datasets, suggesting that it is not influenced by the different normalization and reduction approaches. These results suggest that the PCs are not describing individual trait hotspots, as the top PCs are not associated with the top hotspots, rejecting our hypothesis.

### Evaluating the Impact of PCA on QTL Mapping and Genetic Signal Distribution

One possible reason for the difference in post-hoc summary hotspots and PC vector mapping is genetic linkage. Individual trait QTL mapping accounts for genetic linkage by mapping to the genetic regions controlling each trait. In contrast, by assessing the metabolic variation across genotypes, PCA might identify the genetic linkage associated with phenotypic variation. For example, on chromosome 4, many metabolite QTLs are in linkage with each other, and the PCA may identify vectors that coalesce trait variation across this linkage (Figure 3). In contrast, individual trait mapping will primarily report the marker with the highest significance (Figure 7). This suggests that the use of PCA in QTL mapping may spread the causation associated with a genomic region across multiple PCs. To test this theory, we assessed whether summarizing the percent of total variation linked to each genomic position across all PCs identifies similar regions as found with the post-hoc summary. We calculated the weighted effect of QTLs by multiplying their effect size by the proportion of variance of the PCs with which they are associated. This provides a local estimate of how much total variance is described by a QTL for a PC. We then summed the weighted effect across all QTLs for each genomic position (Figure 6A-6D). Interestingly, this analysis showed that by combining the weighted effects of PCs, the genomic distribution of total variance is comparable to the post-hoc hotspot analysis based on individual traits. For example, the two hotspots on chromosome 2 (Figure 2) are also found as major peaks for total matrix variance (Figure 7). Thus, while PC QTLs ultimately capture the same total variance in the metabolomics data, the PCA creates genetic associations that are not directly linked to specific metabolites.

**Figure 2:**
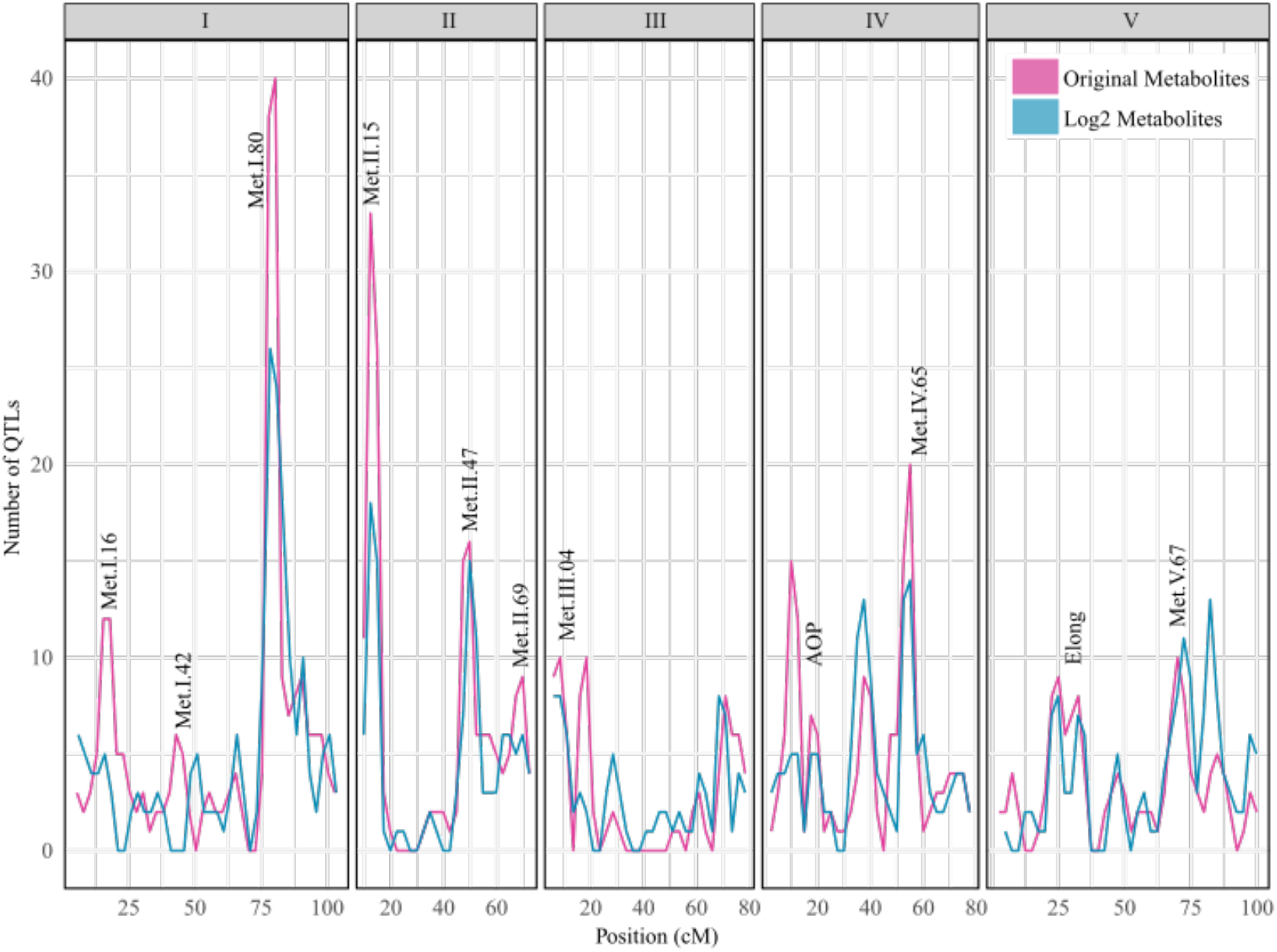
Metabolomic QTL analysis using R/ qtl2. The pink line represents the number of metabolites per 5cM window after mapping unadjusted metabolite data with R/qtl2. The blue line is the mapping with the same metabolomic dataset following a log2 transformation and a 5cM window. Each panel I-V represents an *A. thaliana* chromosome. The metabolite hotspots located by Rowe et al. 2008, are labeled on the QTL plot.

### QTL Hotspots associated with Transcripts

To test if the effects of data reduction on QTL mapping are similar for other omics datasets, we applied the same approaches to a transcriptomics dataset from the same RIL population. We used the expression of 16,838 transcripts measured under two different conditions, experimental (treated with salicylic acid; SA) and control (treated with silwet; SW). This dataset is highly dimensional, with trait variation distinct from the metabolomics dataset. While the metabolite dataset may benefit from normalization using a correlation matrix due to its concentration-based scale, the transcriptomics data is count-based and may benefit more from a log-transformation.

Using the mean expression values for all transcripts, R/qtl2 located 71,862 SA QTLs and 71,678 SW QTLs. These QTLs were distributed across the genome in a pattern similar to that previously observed using the composite interval mapping approach (Figure 4A and Supplemental SLW output tables). After performing log2 transformation to normalize the expression data for each transcript, 72,267 SA QTLs and 72,380 SW QTLs were located (Figure 4B and Supplemental SLW output tables). The log2 transformation did not have a large effect on the number of QTLs detected (Figure 4A and 4B) or the location of the hotspots.

The largest hotspot, located on chromosome 2 at 12.5cM, identified 4,384 SA transcripts in the non-normalized dataset, and 4,350 SA transcripts in the log2-transformed dataset, of which 4,066 transcripts were shared across methods. Similarly, that hotspot hosted respectively 4,100 and 4,055 SW transcripts for the untransformed and log2-transformed data, of which 3,735 transcripts were shared across methods (Figure 4). Thus, the log2 transformation slightly shifted individual transcript QTLs but was consistent in the pattern of hotspots (Figure 4), as also observed in the metabolite dataset.

### PCA Consolidation of Large Transcript Datasets

To test how data reduction affects transcriptomic QTL mapping, correlation and covariance based PCA were performed on both the SA and SW mean expression values, and 211 PCs were used for QTL mapping. The mapping of correlation-based PCs identified 270 QTLs for SA, and 259 QTLs for SW. The mapping of the covariance-based PCs identified 290 QTLs for SA and 271 QTLs for SW (Figure 5, Supplemental Figure 3A to 3D, and Supplemental SLW output tables). Similar to the pattern observed in the metabolite dataset, the PC QTLs were largely evenly distributed across the genome. Furthermore, the first PCs did not identify the post-hoc hotspots. Instead, many QTLs were associated with PCs, and genomic regions were identified across multiple PCs. (Figure 5, Supplemental Figure 3A to 3D, and Supplemental SLW output tables). Comparably, QTLs remained identifiable across the top 100 PCs (Figure 5, Supplemental Figures 1 and 2). As observed in the metabolite dataset, the QTL mapping of PCs is tracking a different impression of how the phenotypic variation is partitioned than the post-hoc QTL analysis.

### Total Weighted Effect Size and Proportion of Variance in QTL Mapping

To investigate if the transcript PCs were coalescing linkage and trait variance, we assessed the total dataset variation explained across the genome. As in the metabolite dataset, the PC QTL mapping identified genomic regions controlling total trait variation that track with the post-hoc summary hotspots. This variance summation approach helped recover the major QTL peaks by associating total variance with marker positions (Figure 7 and 8). These findings highlight how PCA can obscure major QTL peaks by distributing genetic signals across multiple components. However, by aggregating the total weighted effect size across markers, we were able to recover key QTL peaks, demonstrating that the genetic signals were not lost but rather dispersed across PCs. While the recovered QTL map does not perfectly match the post-hoc summary approach, it provides a clearer representation than PC-based QTL mapping alone. This suggests that careful consideration of variance distribution is important when applying dimensionality reduction techniques to genetic data, especially if the reduced dataset is completely different from the unreduced analysis.

## Discussion

When conducting data decomposition on omics data, one goal is to simplify the generation of inference from these large datasets. We originally hypothesized that PCA analysis of large omics datasets would largely track with major hotspots, wherein the genetic variation at the largest hotspot affects the most transcripts or metabolites and could be potentially described by one of the first PCs. However, this was not the case, as the PCs did not simply track with the post-hoc hotspots. Moreover, individual PCs often identified multiple QTLs, and individual genetic loci were identified as influencing multiple QTLs. These QTLs were largely spread out throughout the genome and did not provide an initial impression of the population that readily compared to the post-hoc hotspot analysis summarizing across all individual traits (Figures 3, 5, and 6). Previous work has shown that in *A. thaliana* and other populations, it is possible to use the post-hoc hotspots to generate causal hypotheses. For example, querying the transcripts or metabolites associated with a hotspot can identify the molecular network modulated by that QTL and often the causal gene (West et al., 2007; Rowe et al., 2008; Brachi et al., 2015). While it appears that the PCA methods did efficiently capture variance in the dataset, each approach captured a different image of the variation. As such, the method utilized for analyzing QTL and likely GWA analysis of transcriptomics, metabolomics and/or phenomics should be carefully considered to ensure that the specific question of interest is being addressed.

**Figure 3:**
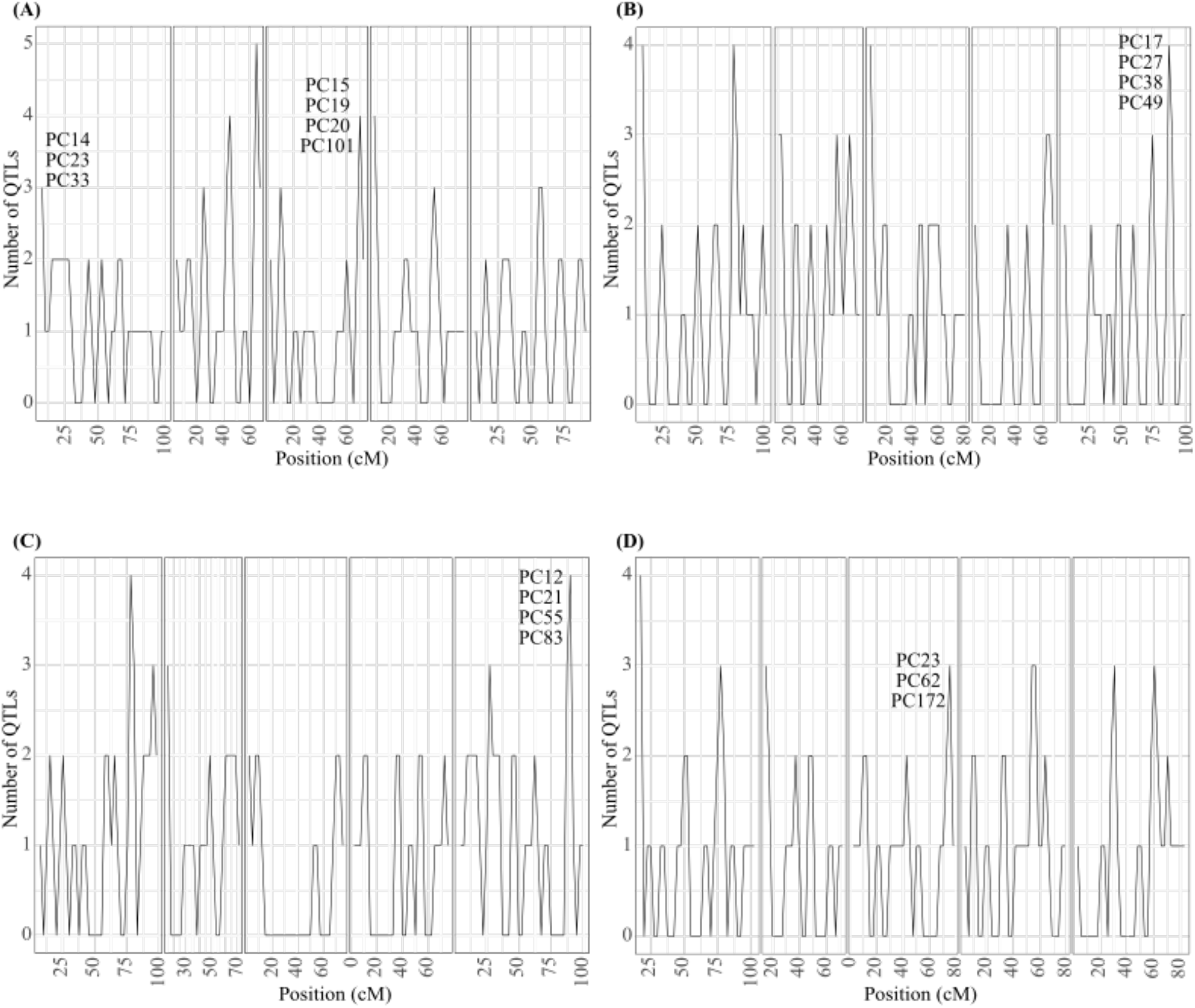
Hotspot analysis of QTL mapping of the 371 metabolites with different data transformation strategies. The QTLs were consolidated within 5cM sliding windows along the five *A. thaliana* chromosomes (each panel along the x-axis). A) QTL mapping of the correlation based PCA on unadjusted metabolites. B) QTL mapping of the covariance based PCA on unadjusted metabolites. C) QTL mapping of the correlation-based PCA on log2-adjusted metabolites. D) QTL mapping of covariance-based PCA on log2-adjusted metabolites. For illustration purposes, some of the larger QTL peaks are labeled with their associated PCs.

**Figure 4:**
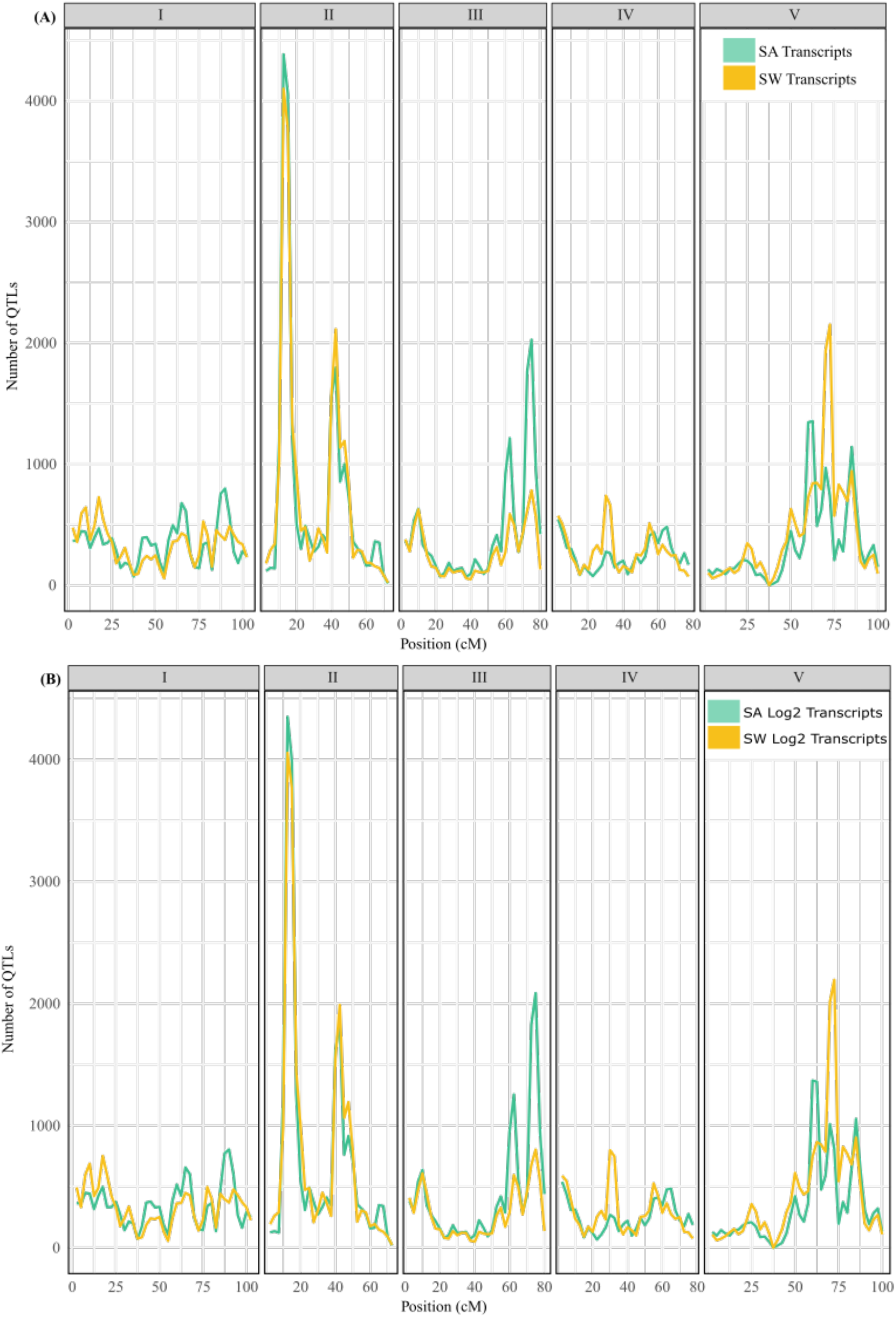
Hotspot analysis of the QTL mapping of 22,810 transcripts for salicylic acid treatment (in green) or silwet control (in yellow) A) without and B) with log2 transformation. Each panel represents the *A. thaliana* chromosomes I-V.

**Figure 5:**
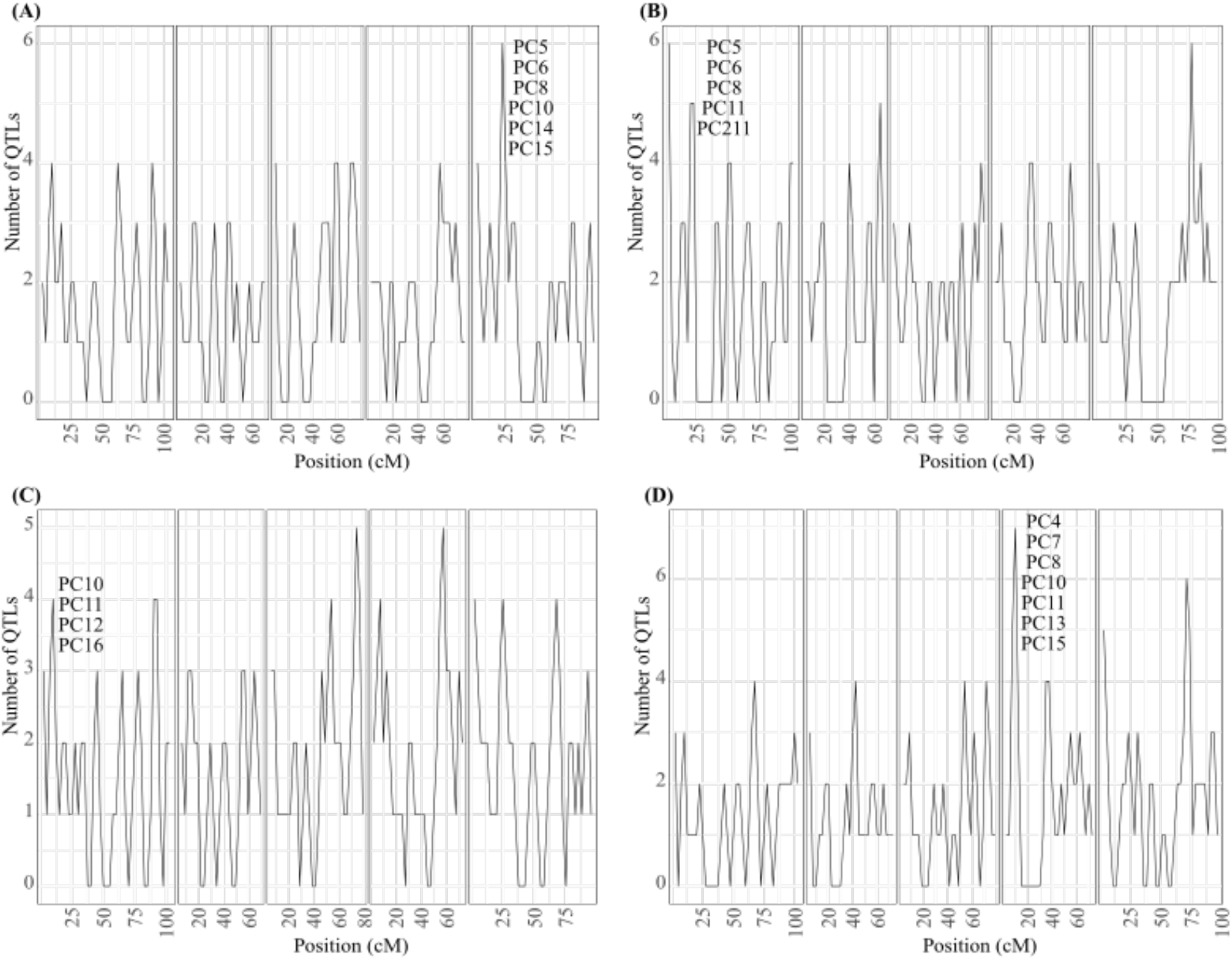
QTL Mapping of 22,810 transcripts after salicylic acid treatment, dimensionally reduced by PCA and analyzed within 5cM sliding windows along the five *A. thaliana* chromosomes (each panel along the x-axis). A) QTL mapping of the correlation-based PCA. B) QTL mapping of the covariance-based PCA. C) QTL mapping of the correlation-based PCA on log2-adjusted gene expression values. D) QTL mapping of the covariance-based PCA on log2-adjusted gene expression values. For illustration purposes, some of the larger QTL peaks are labeled with their associated PCs.

### Genome-Wide Variance Summary

This analysis showed the initial PCA-based mapping flattens the information across the genome in comparison to the post-hoc hotspot analysis. However, this appears to be at least partly generated because the PCs are capturing both trait variation and linkage in the same vector. We tested this by comparing the genomic locations of total variance captured across the PCA of the whole metabolome or transcriptome dataset with those identified in the post-hoc hotspot analysis (Figures 7 and 8). For example, while the individual PCs did not identify the metabolite hotspots on chromosome 2 as particularly important, these two regions explained most of the total trait variation in the metabolite PCA (Figures 3, 5, 6-8). One possible explanation for this disconnect is that the PCA is run on the total dataset prior to QTL mapping and may therefore incorporate aspects of genetic linkage before QTL mapping occurs. For example, if a set of metabolites is influenced by two linked QTLs, the PCA would likely create a vector describing the combined influence of by using the RILs that share linkage between the QTLs. Another PC may then capture the recombinant effects between these QTLs. This would essentially mean that the PCs incorporate genotype/RIL information prior to the QTL mapping by creating population structure. This structuring would not occur when analyzing individual traits and accounts for the observed disconnect between the two approaches. This could potentially create a disconnect between the two approaches. In summary, we show that data reduction needs to be carefully vetted for its usage in genetic mapping to ensure that the specific question of interest is being addressed.

**Figure 6:**
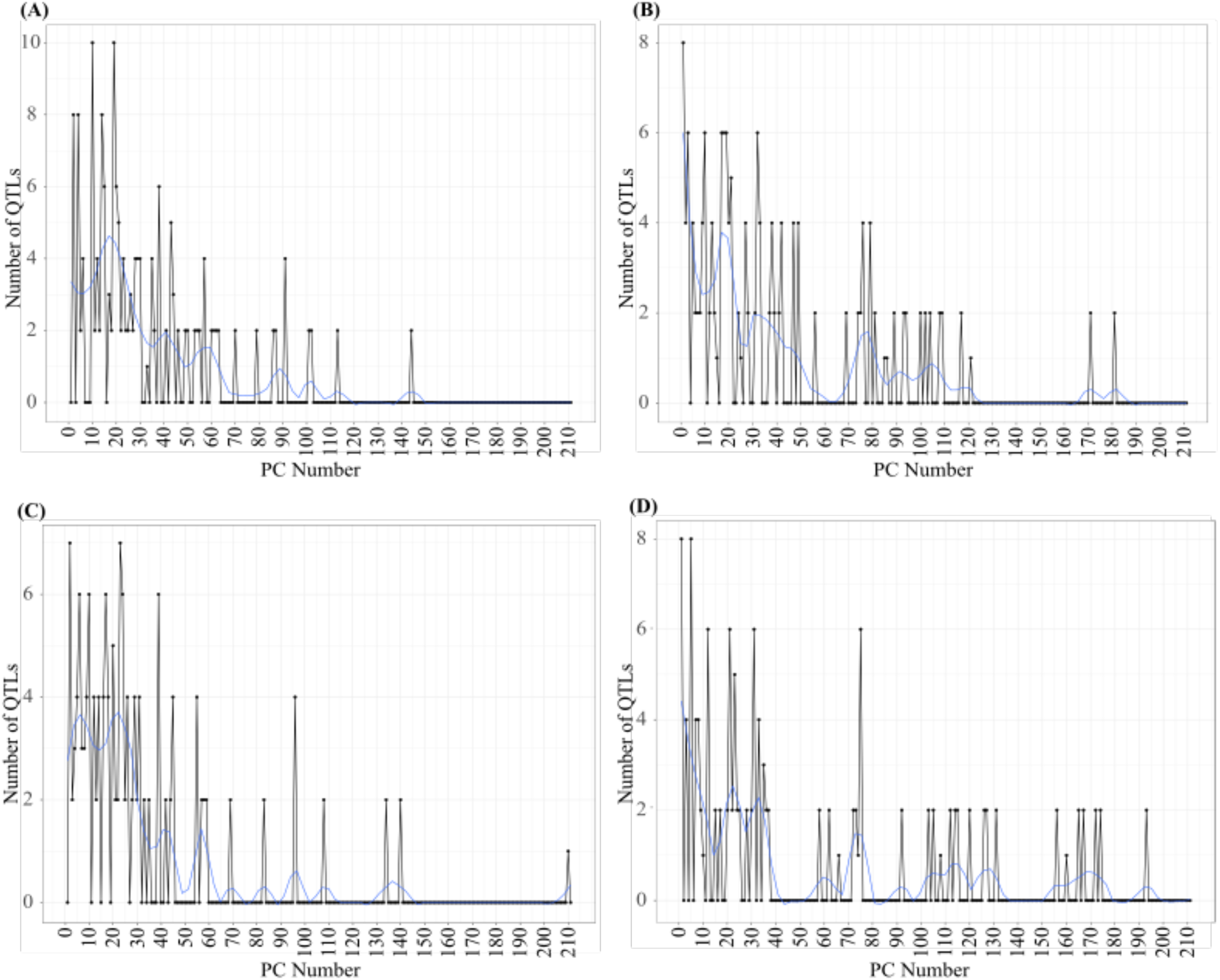
The number of metabolite QTLs associated to each of the 211 principal components. The general trend is plotted in blue with Loess smoothing at 10% and Alpha=0.2. A) Number of metabolite QTLs associated with each correlation-based PCs. B) Number of metabolite QTLs associated with each covariance-based PCs. C) Number of metabolite QTL associated with each log2-transformed correlation-based PCs. D) Number of metabolite QTL associated with each log2-transformed covariance-based PCs.

**Figure 7:**
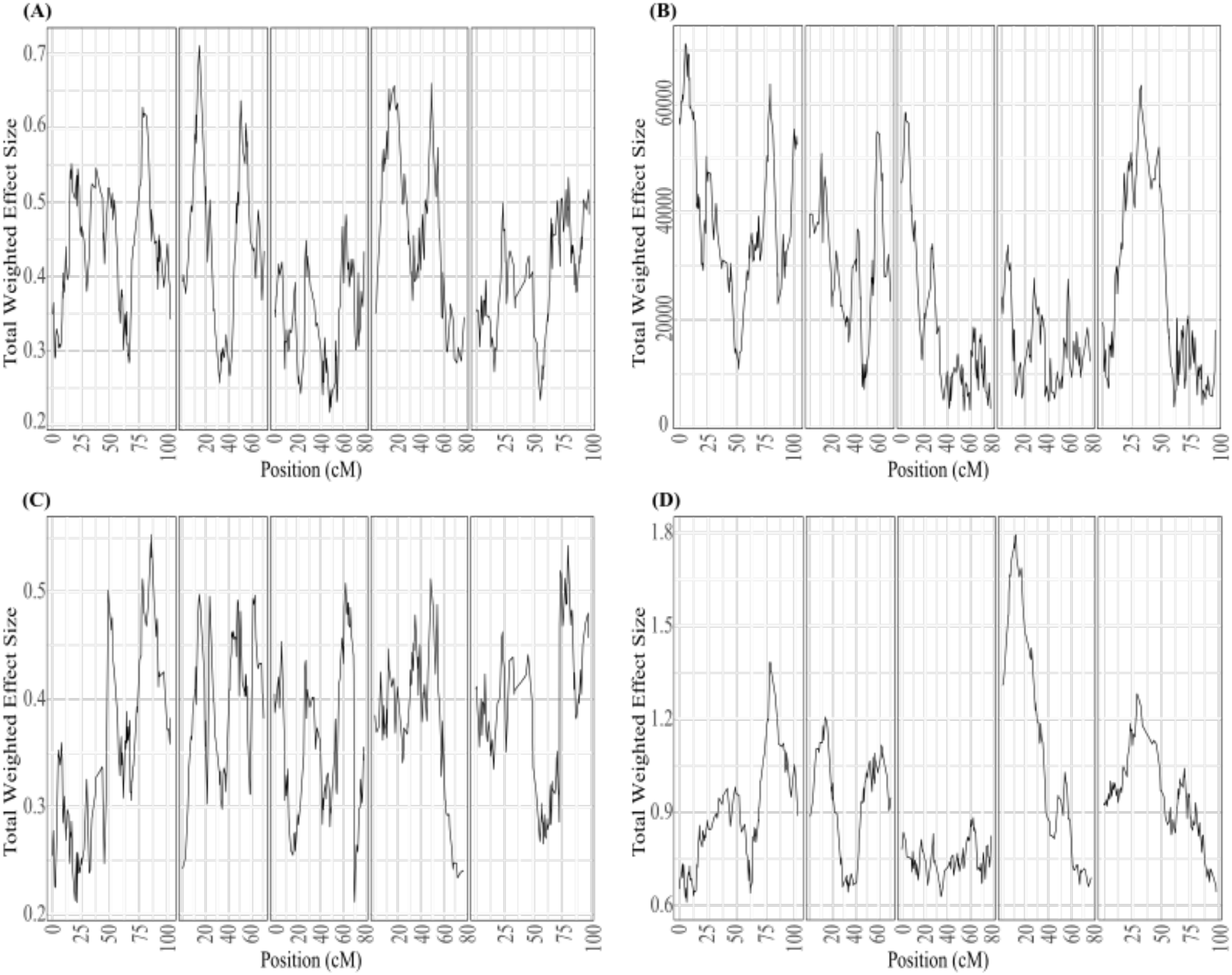
Sum of the total weighted effect per marker in the metabolite data. The PCs’ proportion of variance was multiplied by the effect size to obtain the weighted effect. The weighted effect was summed, and the total weighted effect size for each marker was plotted by position (cM) along chromosomes (I-V). Each panel represents chromosomes 1-5 from left to right. A) Total weighted effect of the PCA correlation matrix. B) Total weighted effect of the PCA covariance matrix. C) Total weighted effect of the PCA correlation matrix with Log2 transformation. D) Total weighted effect of the PCA covariance matrix with Log2 transformation.

**Figure 8:**
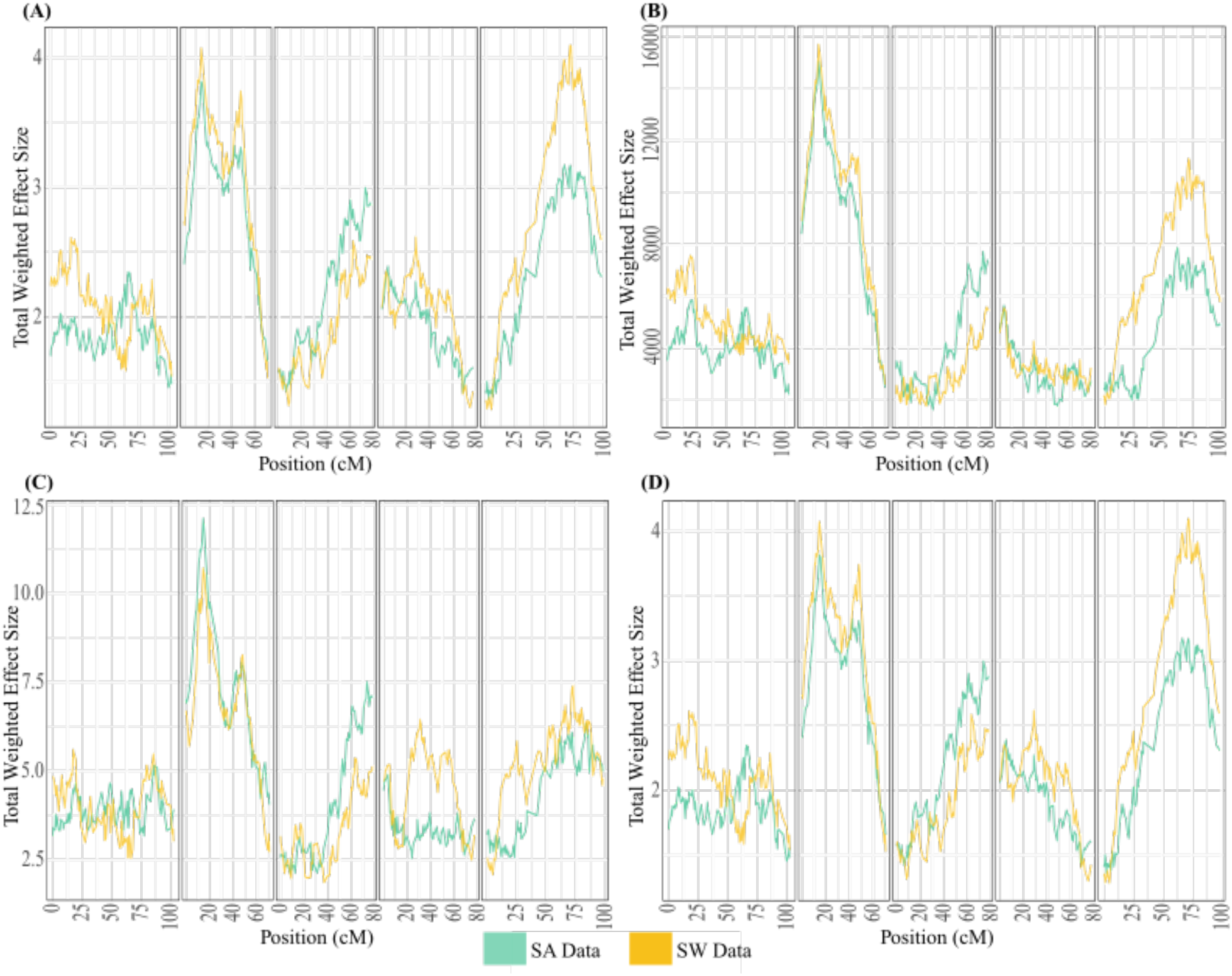
Sum of the total weighted effect per marker in the SA and SW gene expression data. The PCs’ proportion of variance was multiplied by the effect size to obtain the weighted effect. These weighted effect sizes were summed and plotted by position (cM) along chromosomes (I-V). SA transcripts are in green and SW in orange. A) Total weighted effect of the PCA correlation matrix. B) Total weighted effect of the PCA covariance matrix. C) Total weighted effect of Log2-transformed PCA correlation matrix. D) Total weighted effect SA of the Log2-transformed PCA covariance matrix.

## Acknowledgments

We would like to thank Daniel Runcie for extensive discussions about this analysis and the results.

## Funding

Funding for this work was provided by grants to DJK by NSF MCB 1906486 and IOS 2020754.

**Supplemental Figure 1:**
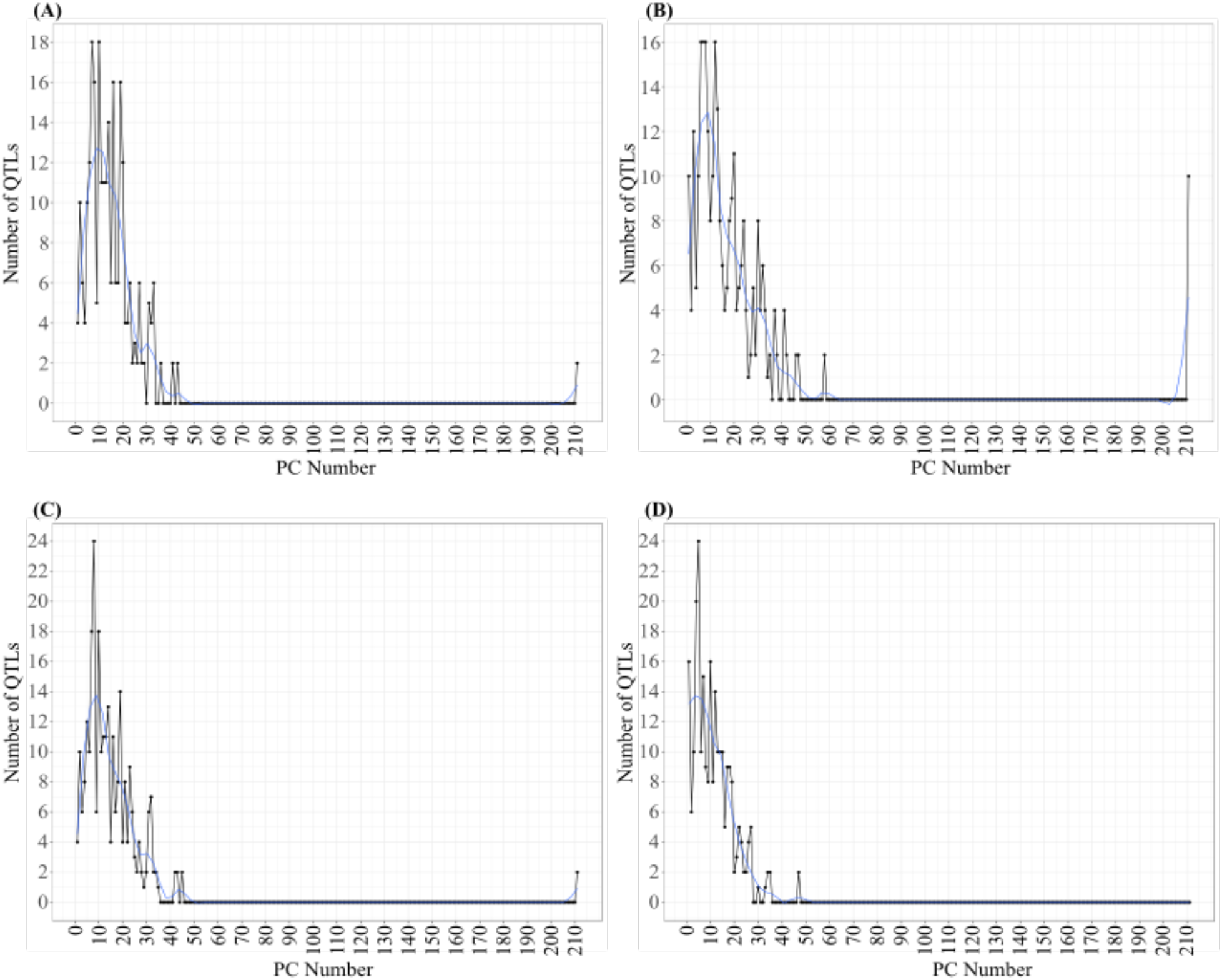
Number of SA QTLs associated with each PC. In blue, Loess smoothing line at 10%, Alpha=0.2. A) PCA correlation matrix. B) Covariance matrix. C): Correlation matrix with Log2 transformation. D) Covariance matrix with Log2 transformation.

**Supplemental Figure 2:**
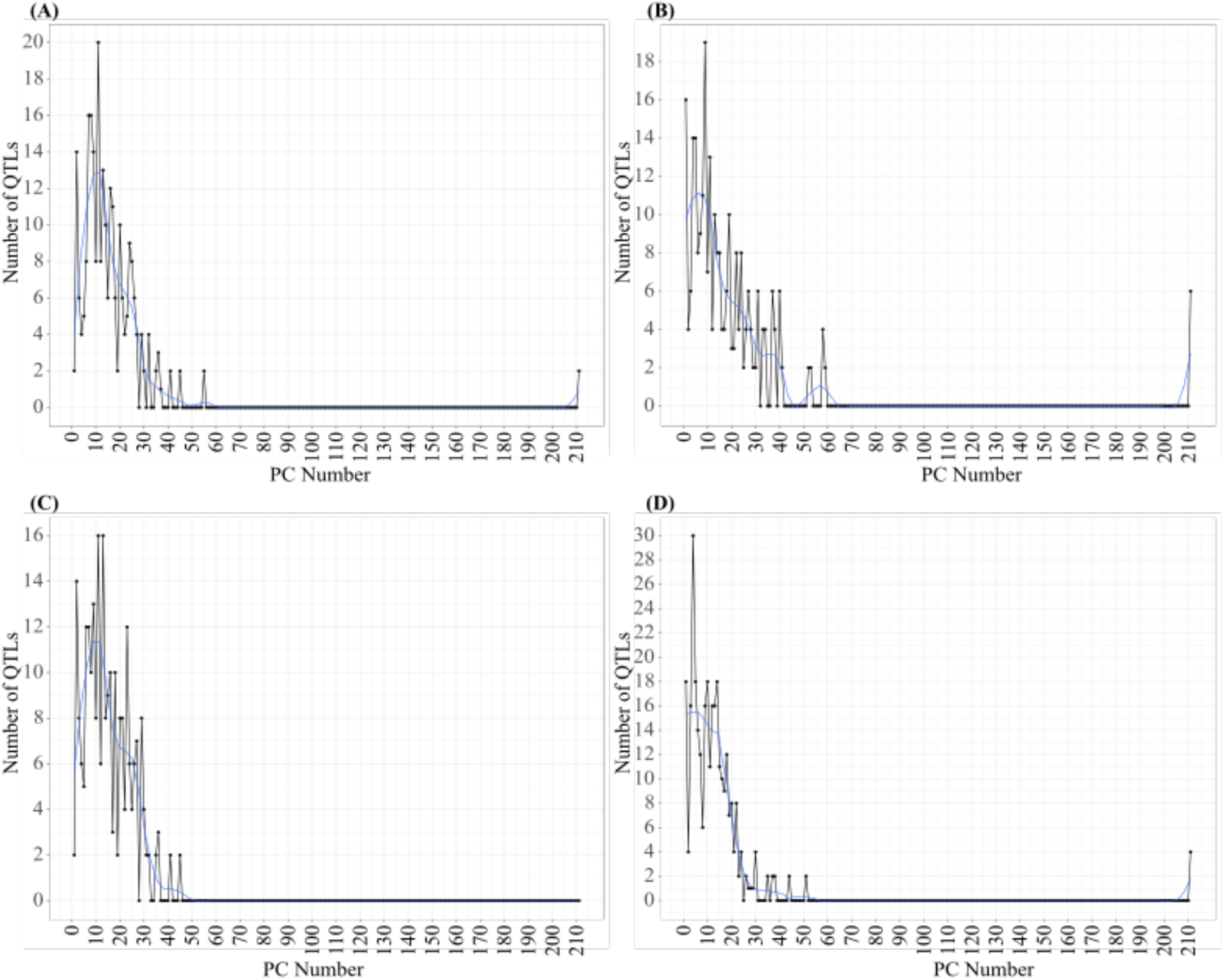
Number of SW QTLs associated to each PC. In blue, Loess smoothing line at 10%, alpha=0.2. A) PCA correlation matrix. B) Covariance matrix. C) Correlation matrix with Log2 Transformation. D) Covariance matrix with Log2 Transformation.

**Supplemental Figure 3:**
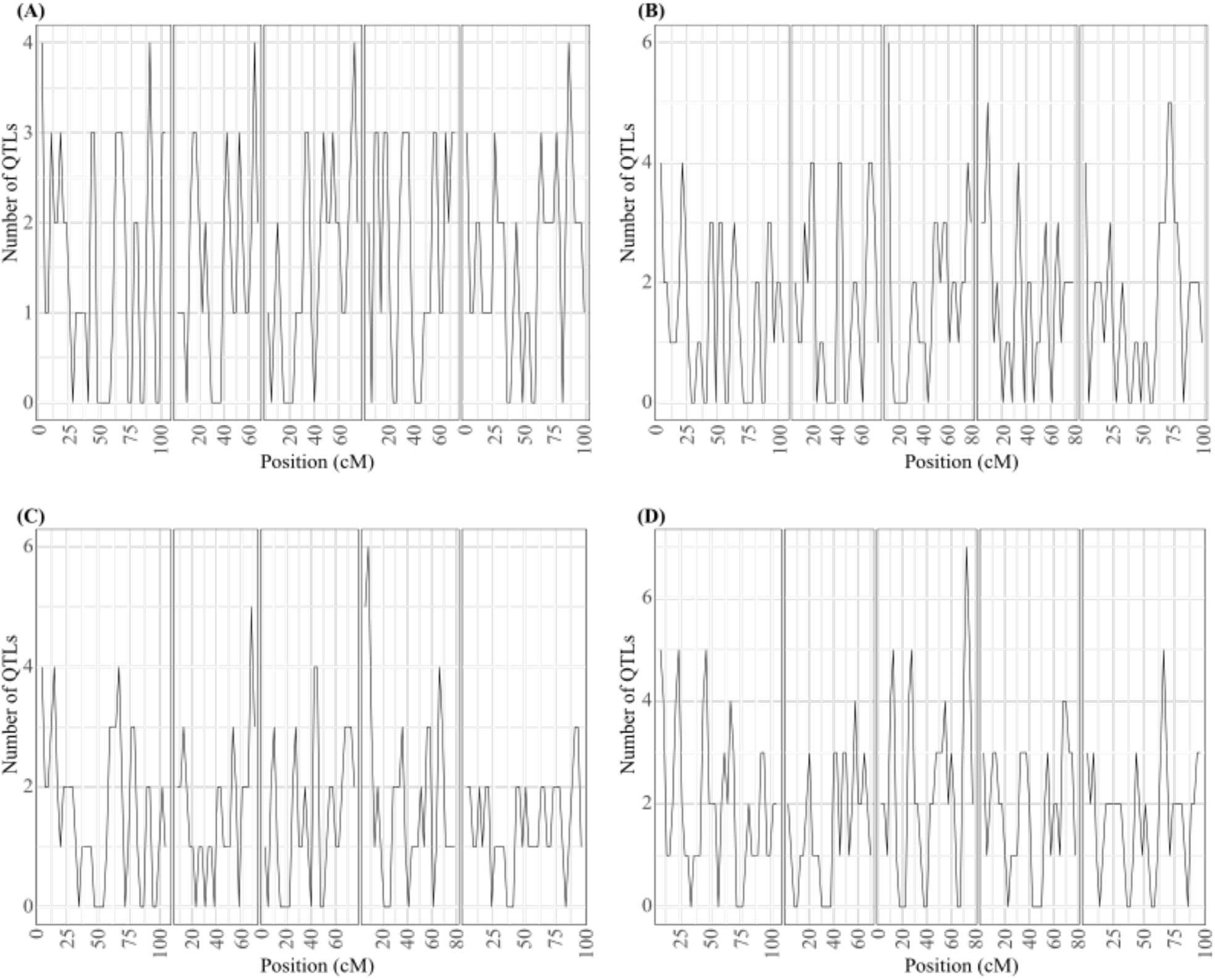
QTL Mapping of 22,810 transcripts after silwet treatment, dimensionally reduced by PCA and analyzed within 5cM sliding windows along the five *A. thaliana* chromosomes (each panel along the x-axis). A) QTL mapping of the correlation-based PCA. B) QTL mapping of the covariance-based PCA. C) QTL mapping of the correlation-based PCA on log2-adjusted gene expression values. D) QTL mapping of the covariance-based PCA on log2-adjusted gene expression values.

## Notes

### Competing Interest Statement

The authors have declared no competing interest.

